# Computational analysis of the *Plasmodiophora brassicae* genome: mitochondrial sequence description and metabolic pathway database design

**DOI:** 10.1101/335406

**Authors:** S. Daval, A. Belcour, K. Gazengel, L. Legrand, J. Gouzy, L. Cottret, L. Lebreton, Y. Aigu, C. Mougel, M.J. Manzanares-Dauleux

**Author notes:** S. Daval and A. Belcour have contributed equally to the work.

## Abstract

*Plasmodiophora brassicae* is an obligate biotrophic pathogenic protist responsible for clubroot, a root gall disease of Brassicaceae species. In addition to the reference genome of the *P. brassicae* European e3 isolate and the draft genomes of Canadian or Chinese isolates, we present the genome of eH, a second European isolate. Refinement of the annotation of the eH genome led to the identification of the mitochondrial genome sequence, which was found to be bigger than that of *Spongospora subterranea*, another plant parasitic Plasmodiophorid phylogenetically related to *P. brassicae*. New pathways were also predicted, such as those for the synthesis of spermidine, a polyamine up-regulated in clubbed regions of roots. A *P. brassicae* pathway genome database was created to facilitate the functional study of metabolic pathways in transcriptomics approaches. These available tools can help in our understanding of the regulation of *P. brassicae* metabolism during infection and in response to diverse constraints.

## 1. Introduction

Clubroot, caused by the obligate telluric biotroph protist *Plasmodiophora brassicae* Wor., is one of the economically most important diseases affecting Brassica crops in the world. The pathogen leads to severe yield losses worldwide (Dixon, 2009). Clubroot development is characterized by the formation of galls on the roots of infected plants. Above-ground symptoms include wilting, stunting, yellowing and premature senescence. Chemical control of the disease is difficult and/or expensive. Agricultural practices, such as crop rotation, and cultivation of resistant Brassica varieties are the main methods for disease control (Diederichsen et al., 2009).

The pathogen has a complex life cycle comprising three stages: survival in soil as spores, root hair infection, and cortical infection. First, resting spores can survive for a long period (up to fifteen years) in the soil. Secondly, germination of haploid resting spores releases primary zoospores which infect root hairs, leading to intracellular haploid primary plasmodia. In the third stage, after nuclear division, secondary zoospores may be released into the soil and penetrate the cortical tissues. The step of released zoospores in the soil before invading the root cortex is not universally accepted among the research community. During this last stage, the pathogen develops into secondary multinucleate diploid plasmodia which cause the hypertrophy and hyperplasia of infected roots into characteristic clubs (Ingram and Tommerup 1972; Bulman and Braselton 2014). Secondary plasmodia finally release a new set of haploid resting spores.

*P. brassicae* is a member of the order Plasmodiophorid within the eukaryote supergroup Rhizaria. This protist taxon includes other important plant pathogens such as *Spongospora subterranea,* causal agent of powdery scab on potato, and *Spongospora nasturtii*, the causal agent of crook root in watercress (Burki et al., 2010).

Several *P. brassicae* genomes are available, including one reference genome from a European isolate (Schwelm et al., 2015), and additional re-sequencing of five Canadian (Rolfe et al., 2016) and one Chinese isolates (Bi et al., 2016). The pathotype classification of these isolates is difficult to compare because they have been characterized using different differential hosts sets. In these previous studies, the main characteristics of the *P. brassicae* genome were described, such as its small size (24.2 - 25.5 Mb) and high gene density. Thirteen chitin synthases were described, suggesting an important role for this gene family in resting spore formation (Schwelm et al., 2015). Secreted proteins, described to play a role in plant pathogenic organisms in suppressing the plant defense responses and modifying the host metabolism, were previously predicted *in silico* in *P. brassicae*, but for most of them, no putative function could be assigned (Schwelm et al., 2015; Rolfe et al., 2016). The *P. brassicae* genome also contains genes potentially involved in host hormone manipulation, such as the auxin-responsive Gretchen Hagen 3, isopentenyl-transferases, methyltransferase and cytokinin oxidase (Schwelm et al., 2015). The clubroot genome also appears to be lacking several metabolic pathways, a characteristic of the eukaryotic biotrophic plant pathogens (Spanu et al., 2010; Baxter et al., 2010; Kemen et al., 2011). The missing genes encode proteins involved in sulfur and nitrogen uptake, as well as the arginine, lysine, thiamine and fatty acid biosynthesis pathways. In addition, only a few carbohydrate active enzymes (CAZymes), involved in the synthesis, metabolism and transport of carbohydrates, were found. Characterization of the *P. brassicae* genome is, however, still incomplete as the above data were from a limited number of genotypes and conditions.

Schwelm et al. (2016) recently summarized the current life stage-specific transcriptomics data related to the molecular events in the life cycle of *P. brassicae*. During spore germination and at the primary zoospore stage, enzymes involved in chitinous cell wall digestion are highly expressed. Several metabolism pathways such as the starch, citrate cycle, pentose phosphate pathway, as well as pyruvate and trehalose metabolism were also very active (Schwelm et al., 2015). Although the molecular mechanisms involved in primary root hair infection are unknown, the secondary phase has been better described. During re-infection by the secondary zoospores in root hairs or the root cortex, genes involved in basal and lipid metabolism were highly expressed (Bi et al., 2016). The plant clubroot symptoms (hyperplasia and hypertrophy of the infected tissues leading to the gall phenotype) were found to result from the deregulation of the host plant primary and secondary metabolism (Ludwig-Müller 2009a; Gravot et al. 2011, 2012; Wagner et al. 2012) as well as from the modification of plant hormone homeostasis (Devos et al., 2006; Siemens et al., 2006; Ludwig-Müller, 2009b; Malinowski et al., 2016) by the pathogen. The main plant hormone pathways modified were cytokinin biosynthesis, auxin homeostasis (Siemens et al., 2006; Ludwig-Müller, 2009b; Schuller et al., 2014), and salicylic acid and jasmonic acid metabolism (Lemarié et al., 2015). However, the mechanisms by which the pathogen may drive plant hormone homeostasis are still not completely deciphered. At the end of the *P. brassicae* life cycle, secondary plasmodia transform into resting spores. In these structures, trehalose could play an important role for their long-term survival capacity (Brodmann et al., 2002). This description of the current transcriptomics data in *P. brassicae* shows that some important steps in *P. brassicae* developmental stages are still not well-characterized at the molecular level.

More information is thus needed, in particular from other clubroot genotypes to improve gene and protein descriptions, thus perhaps allowing the discovery of some of the potentially missing pathways or some specific pathotype features. Therefore, to gain further structural and functional knowledge of the *P. brassica*e genome, the first aim of this study was to generate a de *novo*-sequence from a second European isolate. We report here the genome sequence of the European eH *P. brassicae* isolate belonging to a pathotype widespread in France but different to e3, the reference clubroot isolate. The second aim was to improve genome annotation to resolve, for the first time, the mitochondrial genome of this clubroot pathogen. The third aim was to use this improved annotation to predict new metabolic pathways and develop ClubrootCyc, a metabolic pathway database for *P. brassicae*. This new tool gives researchers access to a repertoire of *P. brassicae* metabolic and transport pathways as well as to an Omics Viewer to analyze expression data, which can be updated with new data.

## 2. Materials and methods

### 2.1. Pathogen material and plant inoculation

The *P. brassicae* isolate used in this study was eH. The “selection” isolate eH (*i.e*., an isolate displaying a phytopathological pattern similar to a single spore isolate) was kindly provided by Dr. J. Siemens (University of Berlin) and was described by Fähling et al. (2003). Isolate eH was characterized for pathogenicity on differential *Brassica napus* cultivars according to the classification proposed by Somé et al. (1996) and corresponded to the pathotype P1.

To ensure genetic homozygosity of the pathogen DNA material, resting *P. brassicae* spores were used. The isolate was propagated on the universal susceptible host Chinese cabbage (*Brassica rapa* ssp *pekinensis* cv. *Granaat*). Clubroots were collected, washed and surface sterilized to avoid host and bacterial contamination. Galls were homogenized in a blender with sterile water and separated by filtration through layers of cheesecloth. The resting spores were then separated by filtration through 100 µm then 55 µM sieves to remove plant cell debris. After centrifugation at 2500 g for 5 min, the pellet was carefully treated with DNase RQ1 for 10 min at 65°C and centrifuged at 2500 g for 5 min at 4°C (Manzanares-Dauleux et al., 2000).

### 2.2. DNA and RNA extraction

Prior to nucleic acid extraction for genome sequencing, the purified resting spores were ground in liquid nitrogen using a mortar and a pestle. DNA was purified using the NucleoSpin Plant II Kit (Masherey-Nagel) following the manufacturer’s instructions. The DNA quality was verified on an agarose gel and the quantity was estimated with a Nanodrop 2000 (Thermoscientific). The absence of host DNA plant contamination was tested by PCR using Brassica specific cruciferin primers (forward primer 5’-ggccagggtttccgtgat-3’ and reverse primer 5’-ccgtcgttgtagaaccattgg-3’).

Total RNA was also obtained from purified, non-germinating, resting spores using TRIzol reagent (Invitrogen) following the manufacturer’s instructions. RNA purity and quality were assessed with a Bioanalyser 2100 (Agilent) and quantified with a Nanodrop (Agilent).

### 2.3. Genome sequencing and assembly

For *de novo* sequencing of the eH genome, different libraries were sequenced using 40 µg DNA with Illumina HiSeq 2500 technology (Genoscreen, Lille, France). Three libraries were sequenced: a paired-end shotgun library with an insert size of 430 bp, a 3 kb insert mate-pair Illumina library, and a 5 kb insert mate-pair Illumina library. Paired-end reads were cleaned (N-trimming) and reads shorter than 50 bp were removed. The eH genome was *de novo* assembled (86 bases k-mer) using SOAPdenovo. Gaps were trimmed and gap-closed using GapCloser, scaffolds were filtered (500 nt in size) and redundancy was removed.

For scaffold #43, identified as the mitochondrial genome, assembly was performed with appropriate software, such as MITObim (Hahn et al., 2013), NOVOPlasty (Dierckxsens et al., 2017) and MindTheGap (Rizk et al., 2014).

### 2.4. Transcriptome sequencing and functional annotation

RNA from eH non-germinating resting spores was used for RNA sequencing using 1×100 bp Illumina HiSeq2500. The oriented single end reads were trimmed and cleaned. All non-redundant generated sequences were processed using Velvet for transcriptome assembling and Gmap for alignment to the eH genome sequence. Genome annotation was conducted using EuGene prediction software (Foissac et al., 2008) trained on *P. brassicae* data. The annotation pipeline included in EuGene integrates various sources of evidence, such as high throughput strand-specific RNA-Seq data, intrinsic information provided by coding potential (Interpolated Markov Models), stop and start codon analysis (using a dedicated RBS alignment tool), similarities with known proteins (SwissProt), high quality CDS predictions and ncRNA predictions (tRNAscan-SE, RNAmmer and Rfam-scan softwares). Putative transcript function and Gene Ontology (GO) analyses were performed using Blast2GO (Conesa et al., 2005) with default settings. For more detailed genome annotation and to retrieve information (Enzyme Commission numbers (EC), GO terms, InterPro and pathways), several external databases were interrogated: Reactome (Fabregat et al., 2016), KEGG (Kanehisa et al., 2017), MetaCyc (Caspi et al., 2016), EupathDB (Aurrecoechea et al., 2017), Uniprot (The UniProt Consortium, 2017), Gene Ontology (Ashburner et al., 2000; The Gene Ontology Consortium, 2017), and ECDomainMiner to link known Pfam domains to EC numbers (Alborzi et al., 2017). Carbohydrate-active enzymes (CAZymes) were predicted using the dbCAN pipeline (Yin et al., 2012). SignalP (Petersen et al., 2011) was used to detect secretion signals in protein sequences. The protein sequences predicted to have a secretion signal by SignalP, were then subjected to transmembrane helix prediction with TMHMM (Krogh et al., 2001). Blastn and Blastp were used to search for already known sequences (Altschul et al., 1997). This was done for the *P. brassicae:* i) serine protease (Feng et al., 2010), ii) methyltransferase (Ludwig-Müller et al., 2015), iii) genes detected by dot plot and real time PCR in the Led09 isolate (Feng et al., 2013), iv) e3 genome (Schwelm et al., 2015), v) genes associated with lipid droplets (Bi et al., 2016), and vi) the mitochondrial genome of *S. subterranea* (Gutiérrez et al., 2014). To visualize the data, circular representations of scaffolds were created using Circos (Krzywinski et al., 2009).

Gene annotation of scaffold #43 was performed with the automated MFannot tool (Burger et al., 2013) that predicts group I and group II introns, tRNAs, RNase P-RNA, and 5S ribosomal RNA (rRNA). The annotation was verified with Blastn and Blastp by quering the NCBI databases.

### 2.5. Metabolic pathway database

A GenBank file was created containing detailed DNA sequence information and used to run Pathway Tools version 21 (Karp et al., 2011). This allowed a Pathway Genome DataBase (PGDB) to be created for *P. brassicae*: ClubrootCyc. Pathways were predicted with MetaCyc database version 20.5 (Caspi et al., 2016). PGDB functionalities were validated using data from the transcriptomics analysis of lipid droplets by Bi et al. (2016).

### 2.6. Data Availability

The eH *P. brassicae* genome sequence has been deposited in the public domain in GenBank Data Librairies under accessions SRR6395507, SRR6395506 and SRR6395505 for raw data sequences, and POCA00000000 for the genomic data. The Pathway Genome DataBase ClubrootCyc is available online at https://pathway-tools.toulouse.inra.fr/CLUBROOT

## 3. Results and discussion

### 3.1. Genome sequencing and assembly

For a better understanding of the pathogenesis and metabolic potential of *P. brassicae*, we generated a *de novo* assembled genome of the eH isolate (belonging to pathotype 1 according to Somé et al., 1996). A paired-end shotgun library consisting of 73,082,688 reads (read length of 101 bp and insert size of 430 bp) and approximately 225-fold genome sequence coverage was obtained. Two additional paired-end libraries were also generated. A 3 kb insert mate-pair Illumina library resulted in 97,629,456 reads (read length of 101 bp and insert size of 3200 bp) and a 5 kb insert mate-pair Illumina library resulted in 81,813,222 reads (read length of 101 bp and insert size of 5050 bp). Both libraries displayed also an approximately 225-fold genome sequence coverage. The final assembly consisted of 136 gap-filled scaffolds with a N50 of 741,293 bp and the total sequence length was 24,559,637 bp with a 59.29% GC content. Basic information about the assembled genome and predicted genes is shown in Table 1. The main features of the eH genome were similar to the reference e3 genome, particularly concerning GC content (Figure 1), but the eH genome displayed a less fragmented assembly and a higher number of predicted genes (12,591 in eH and 9,730 in e3). Figure 1 shows a visual representation of the eH genome’s first 43 scaffolds (which represent 95% of the total genome sequence) compared to the other available *P. brassicae* sequences (the sequences published by Rolfe et al. (2016) were not included because data are not available). The gene density, calculated as the number of genes every 20 kb, was nine on average. This gene density is relatively high as previously described (Schwelm et al., 2015; Rolfe et al., 2016), and contributes to the small genome size compared to the two known genomes of free-living Rhizaria, *Bigelowiella natans* (∼100 Mb) (Curtis et al., 2012) and *Reticulomyxa filosa* (∼320 Mb) (Glöckner et al., 2014).

**Table 1.**
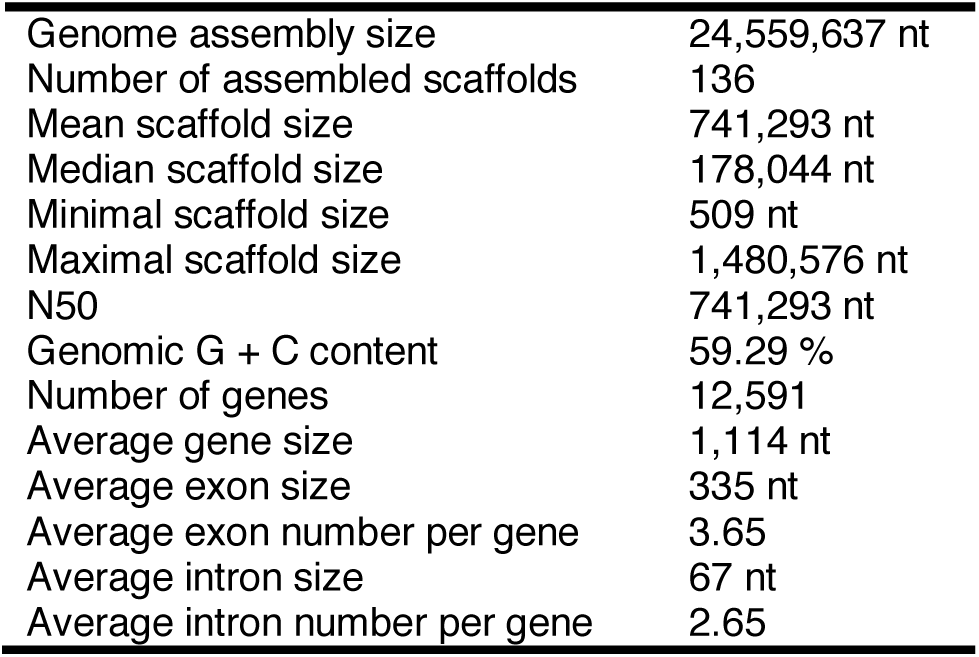
Genome statistics of eH isolate.

**Figure 1.**
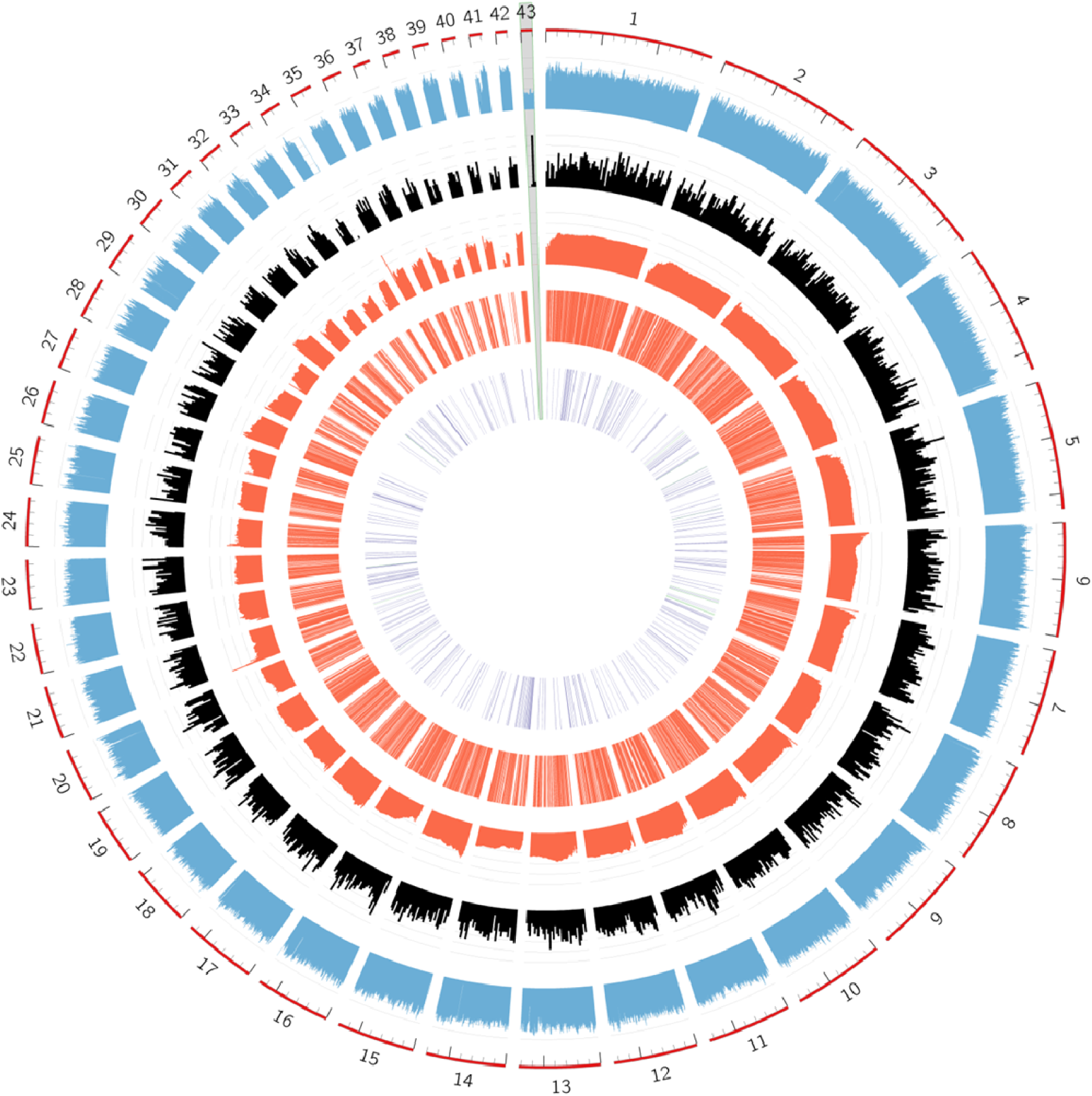
Visual representation of the eH genome first 43 scaffolds. In the first outer circle (A) are represen ted the first 43 scaffolds of the assembled eH genome that correspond to 95% of the total genome sequence. In the second and third outer circles (B and C), the percentage in GC and the gene density (calculated as the number of genes fMJry 20kb) are indicated, respectively. In the two following inner circles, is represented the gene similarity between the isolates eH and e3 (Schwelm et al., 2015): D circle pictures the relative rate every 10kb of the eH genes number to the eH genes number that have similarity to e3 genes (80% threshold); E circle displays this similarity as lines when genes from eH and e3 have 80% similarity. The inner circle shows the gene similarity between the isolates eH end ZJ-1 (E, purple lines; Bi et el., 2016), and eH and Led09 (E, green lines; Feng et al., 2013). The grey scaffold 43 represented the mitochondrial genome.

Gene similarity between the eH and e3 isolates was studied by comparing the gene sequences of both genomes. Using a stringent query span of 80% (meaning that the percentage of alignment coverage of the query sequence should be at least 80%, making sure that we compared genes with common regions), 60% of the genes found in the eH genome showed similarity with the e3 genes. Among the 6,654 alignments displaying the query span of 80%, 4,702 were with 100% gene identity and only 52 were with <60% gene identity (value often used to define genes as highly divergent). For the ZJ-1 isolate’s genome (Bi et al., 2016), annotated regions are not available which made it unfeasible to carry out such accurate comparisons between genes and only contigs were compared which include non-coding regions.

The scaffolds #44 to #242 (corresponding to 5% of the total genome) and in particular scaffolds #88 to #242 were more difficult to assemble (Figure S1) and thus few genes were annotated in these regions.

Scaffold #43 had a lower GC content and corresponded to the mitochondrial sequence (see 3.3.).

### 3.2. Genome annotation

The RNAseq of resting spores, used for annotation, resulted in 23,540,954 reads. The genome annotation was improved compared to the e3 reference genome by interrogating several external databases as described in the materials and methods. The annotation data predicted 12,591 protein-coding genes in eH genome (Table 1). In total, 7,420 genes were associated with GO terms, 4,379 with EC numbers and 8,081 with InterPro signatures. In rare cases of conflicting annotation, the results from manual annotation and Blast2Go were favored.

Carbohydrate-active enzyme sequences of both the e3 and eH genomes were compared using dbCAN, a web server and DataBase for automated Carbohydrate-active enzyme ANnotation. For the eH isolate, 287 enzymes were predicted (Figure S2; Table S1): 89 Glycoside Hydrolases (GH), 111 Glycosyltransferases, 47 Carbohydrate Esterases (CE), 7 Polysaccharide Lyases, 23 Carbohydrate-Binding Modules (CBM), and 10 Auxiliary Activities. GH18, CE4 and CBM18 were the most abundant enzymes (respectively 19, 21, 7 were predicted) within some of these groups. Amongst these 287 eH genes with predicted carbohydrate-active enzyme activity, 223 matched with e3 genes. The list of CAZymes was similar in both the eH and e3 genomes.

Using SignalP and TMHMM to predict the secretome, 741 protein sequences were found to harbor a secretion signal prediction without transmembrane helix. Furthermore, 543 sequences corresponded to small proteins (less than 450 amino acids). Schwelm et al. (2015) analyzed the e3 genome and predicted 553 secreted proteins including 416 with a length smaller than 450 amino acids. Among the proteins with a signal prediction, 355 were common to e3 and eH genomes (Table S2). The higher number of predicted proteins found in the eH genome could be due to the annotation using Eugene which probably created more gene sequences compatible with secretome pattern detection. In addition, the use of updated software versions may have also upgraded the secretome.

An additional annotation was carried out by blasting eH data against clubroot published sequences. All available sequence data for concerning eH and other *P. brassicae* isolates are summarized in Table S3. Ninety-nine percent and 100% identity matches were observed for the *P. brassicae* methyltransferase gene (Ludwig-Müller et al., 2015) and the serine protease gene (Feng et al., 2010), respectively. Three hundred and eighty eight out of the 426 genes described by Bi et al. (2016) involved in lipid metabolism were found in the eH genome (Table S3B). From the 118 genes identified by Feng et al. (2013) in the Led09 isolate’s genome, 23 matched the eH genome (Table S3C). Genes identified as being involved in pathogenicity (*i.e.* with functions such as P-type ATPase activity, Mps1 binder-like protein) were mapped to the eH genome. To complete the annotation, out of the 9,732 genes in the e3 genome, 6,626 were matched to the eH genome (Table S3A). By exploiting this nucleotide variability, additional genome sequencing of *P. brassicae* isolate may allow more precise diagnostic molecular tools than the current pathotyping systems to be developed.

These annotations were added to the corresponding eH genes using python scripts querying the Uniprot database. Using this updated annotation, the genes were classified into subsystems whose main terms are described in Figure 2. Most of the annotated genes were classed in the protein-binding (1,433) GO term category. This class, not accurately defined for their function, still remained high because of the lack of functional data in Plasmodiophorid genomics resources. Ninety-six genes encoding potential RNA-binding proteins having at least one RNA-binding domain were identified. This protein family could play a role in *P. brassicae* pathogenicity, as described in *Ustilago maydis*, the basidiomycete causative agent of corn smut disease (Becht et al., 2005). From the InterPro (IP) database, the ankyrin repeat (258 and 259) was found to be the major protein domain. The ankyrin repeat is the most common protein-protein interaction motif in nature. It is predominantly found in eukaryotic proteins, but also in many bacterial pathogens which employ various types of secretion systems to deliver ankyrin-containing proteins into eukaryotic cells where they mimic or manipulate various host functions (Al-Khodor et al., 2010). In eukaryotes, proteins with ankyrin repeats are involved in many cellular functions, such as transcriptional regulation, signal transduction, cell cycle but also in virulence in Amoebae, suggesting an adaptation to their intracellular localization (Molmeret et al., 2005). The high number of ankyrin repeats in the *P. brassicae* genome could suggest a role in its capacity to invade host plant cortical cells.

**Figure 2.**
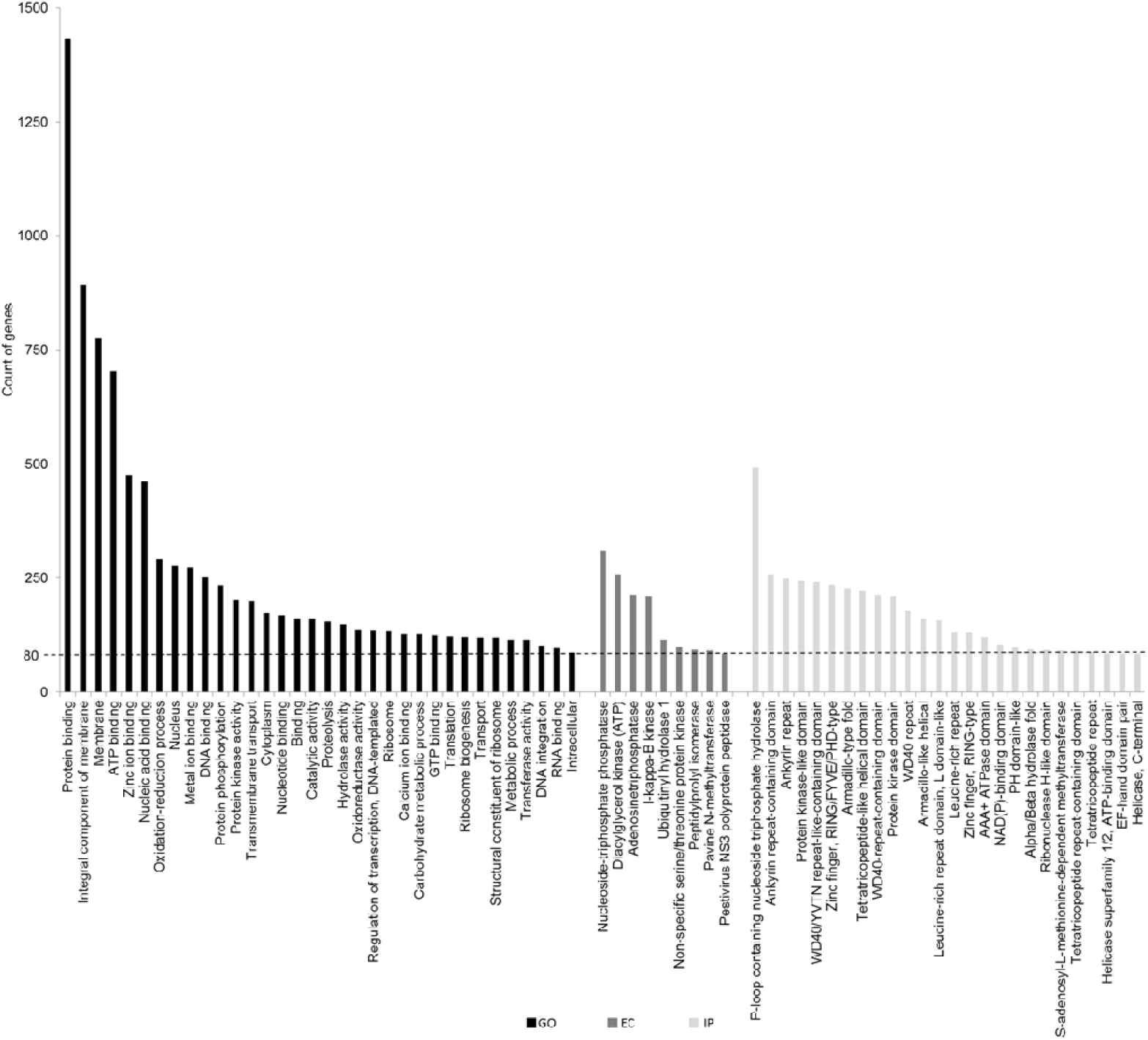
Assignment of genes in different annotation categories. GO, Gene Ontology term (Blast2GO); EC, Enzyme Commission number(Blast2GO and ECDomainMiner); IP, lnterpro term (Blast2GO and Gene Ontology Database). The terms in which at least BO genes were classified are represented.

To summarize the long list of GO terms by removing redundancy, a semantic similarity-based graph was constructed with REViGO (Supek et al., 2011) (Figure S3). This allowed a representative subset of the terms to be found using a clustering algorithm that relies on semantic similarity measures. The Revigo Treemap showed a high representation of the GO terms linked to protein phosphorylation. Changes in the phosphorylation state have a broad impact on cells because they affect many processes such as the subcellular protein localization, protein-protein interactions or signaling cascades (Johnson, 2009). Moreover, interference of the host phosphorylation machinery is a common strategy used by pathogens to promote infection and development in host tissues (Albataineh and Kadosh, 2016), particularly in plant-pathogenic fungi (Turra et al., 2014). A better understanding of the regulation of protein phosphorylation could help in developing targets for regulation of *P. brassicae* infection processes. Another GO term found frequently in the *P. brassicae* genome annotation was related to lipid metabolism. This is consistent with Bi et al. (2016) who reported proteins associated with lipid droplets in resting spores of *P. brassicae* and showed that lipid conversion is high in this protist during cortex infection.

### 3.3. Mitochondrial genome sequence

In the circular genome representation (Figure 1), scaffold #43 showed a low GC content and one match with a *P. brassicae* mitochondrial gene (Siemens 2002, Unpublished, NCBI ID: AF537102). In addition, as a high AT content was found in some Plasmodiophorid mitochondrial genomes, for example in *S. subterranea* (Gutiérrez et al., 2014), these observations suggested that the scaffold #43 could correspond to the *P. brassicae* mitochondrial genome. To ensure that no errors occurred in its assembly, a new assembly of this scaffold #43 was performed using software specifically designed for organelle assembly (NOVOPlasty). Both assemblies were similar suggesting consistent results.

The newly assembled sequence of eH scaffold #43 is 102,962 bp, with a 24.67% GC content (Figure 3) and is very similar (Max score = 91831, Total score = 1.728e+05, Query cover = 90%, E-value = 0.0, Identity = 99%) to scaffold #49 of the genome of the *P. brassicae* isolate ZJ-1 (Bi et al., 2016). The size of the mitochondrial *P. brassicae* sequence (102,962 bp) is larger than that of *S. subterranea* (37,699 bp) (Gutiérrez et al., 2014). Such diversity in genome size was previously described in Chlorarachniophytes, a small group of algae from the supergroup Rhizaria phylum Cercozoa, in which the mitochondrial genomes ranged from 34 to 180 kbp (Tanifuji et al., 2016). Genome sizes vary considerably because of variation in the intergenic regions and intron content, which both contribute to mitochondrial genome plasticity. In *P. brassicae*, numerous long introns were found in mitochondrial genes in contrast to the *Lotharella oceanica* (Rhizaria group) and *B. natans* mtDNAs (Tanifuji et al., 2016).

**Figure 3.**
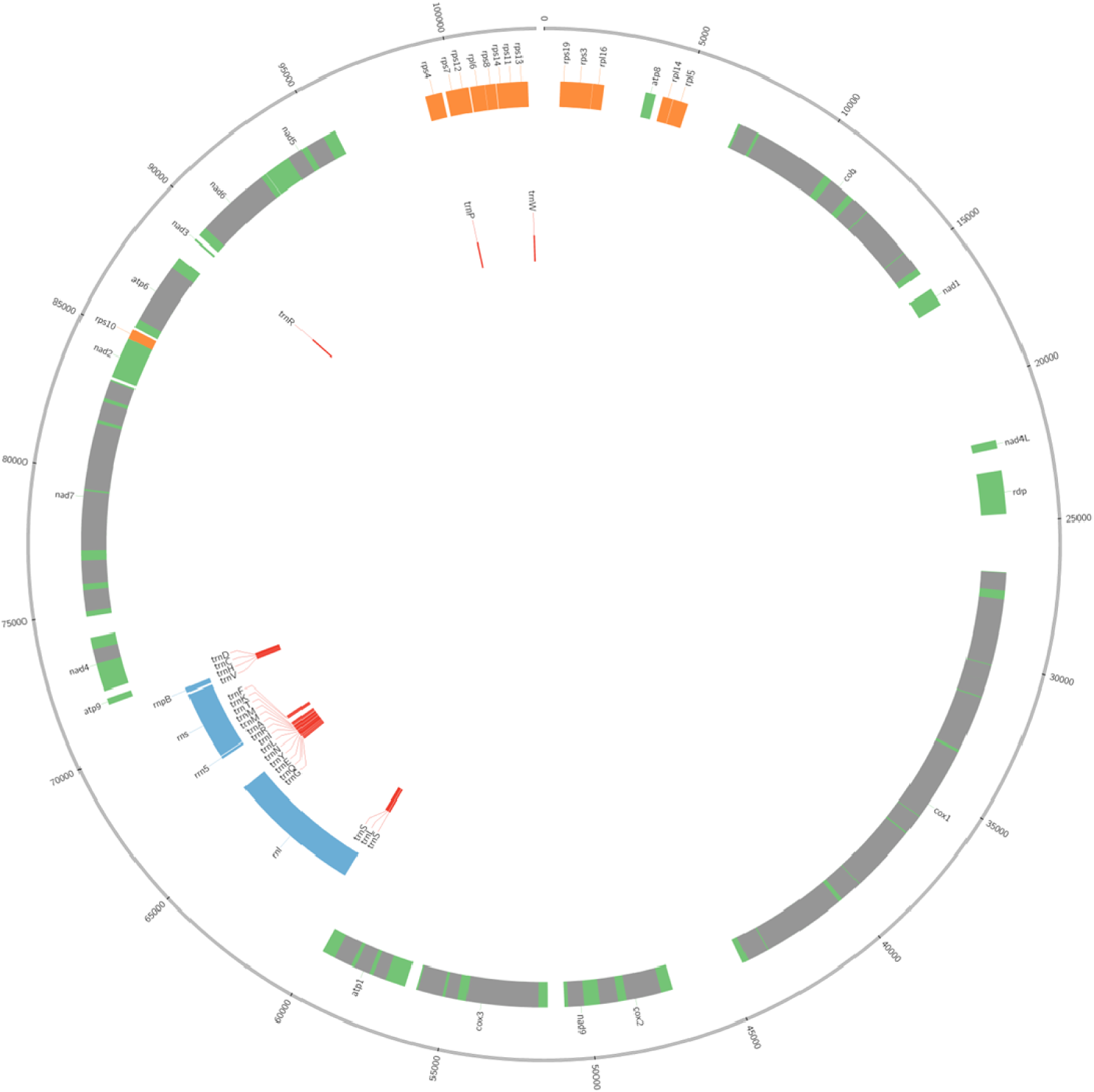
Circular DNA map of the mitochondrial genome of *Plasmodiophorabrassicae.* The outer circle shows the position of protein coding genes with axons represented in green and introns in grey, and the position of ribosomal potein coding genes in orange. The inner rings depicts the position of structural RNAs: rRNA and one ncRNA (rnpB) in blue, and tRNA in red.

Interestingly, despite the significant difference in total size, the *P. brassicae* and *S. subterranea* mitochondrial genomes exhibit similar architecture and high synteny. Both mitochondrial sequences are indeed similar: when the blast best-hit analyses of the *P. brassicae* mitochondrial genes were displayed for the “protein sequences” (32 protein-coding genes (Table S4)) and for the “complete gene sequences” (the 28 other types of genes (Table S5)), 60 genes were found among which 56 were also found among the 59 described in *S. subterranea* (Gutiérrez et al., 2014). Genes coding for proteins involved in respiration and oxidative phosphorylation (17 genes: atp 1-7, 4L, 9, cob, cox 1-3, atp 1, 6, 8, 9), and for ribosomal subunit proteins (10 for small subunit: rps3, 4, 7, 8, 10, 11, 12, 13, 14, 19 and 4 for large subunit: rpl 5, 6, 14, 16) were identified. As in *S. subterranea,* 24 genes coding for tRNA were found in the *P. brassicae*, but trnH described in *S. subterranea* was not found in *P. brassicae* and tRNQ was found only in *P. brassicae* (and not in *S. subterranea*). Interestingly, the protein-coding genes in *P. brassicae* displayed a mosaic structure with small parts spread across the scaffold, as described for mitochondrial sequences of the chlorarachniophyte *B. natans* (belonging to the Rhizaria group) and the model cryptophyte protist *Guillardia theta*; both algal nuclear genomes are mosaics of genes derived from the host and endosymbionts (Curtis et al., 2012). No transposable elements were detected in the *P. brassicae* mitochondrial genome.

### 3.4. Metabolic pathway prediction

Pathway Tools Software was used to create a *P. brassicae* specific database. Taking advantage of the improved annotation described here and available databases (Uniprot, Gene Ontology and ECdomainMiner), a Genbank entry was created. Using this file as input, the PathoLogic component of Pathway Tools could be used to create a new Pathway Genome DataBase (PGDB) containing the predicted *P. brassicae* metabolic pathways. The database is called ClubrootCyc (Figures 4 and S4) and predicts 115 pathways, 1,938 enzymatic reactions, 71 transport reactions, 4,468 enzymes, 123 transporters, and 1,567 compounds (https://pathway-tools.toulouse.inra.fr/CLUBROOT). It links genomic data to additional information such as protein annotation, enzymatic reactions and pathways.

**Figure 4.**
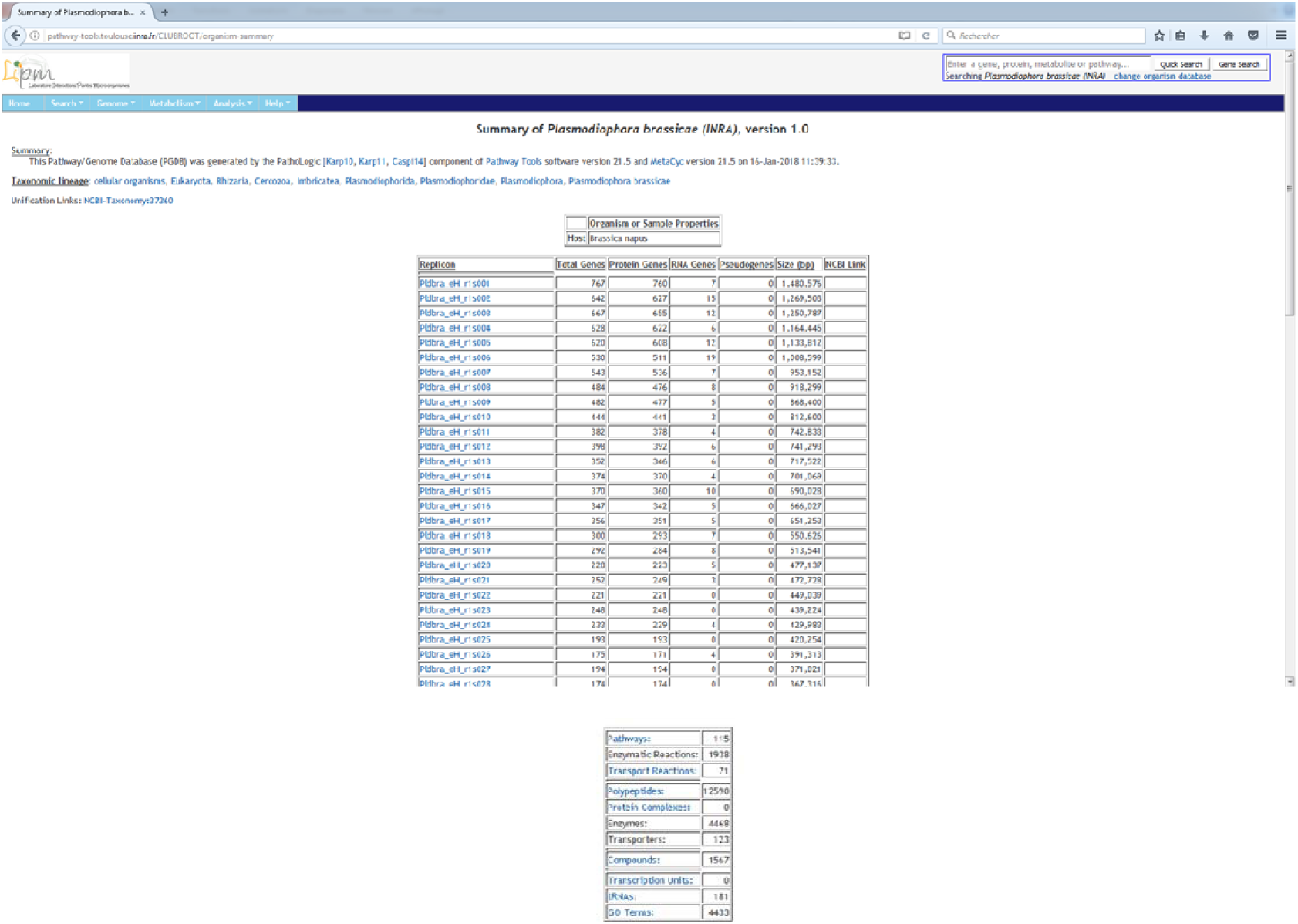
A view of ClubrootCyc homepage. Its mains features and functionalities, such as the structure of each scaffold, the gene local context, the pathway genome overview, and the toolbar for several searchs are displayed in the Figure S4. The complete pathway genome database is available at http://pathway-tools.toulouse.inra.frforganism-summray?object=CLUBROOT

To validate the relevance and functionality of the ClubrootCyc database, the metabolic reconstruction was tested with available published transcriptomics data described by Bi et al. (2016). Visual analysis of omics gene-expression, highlighted in Figure S5, showed that the major pathway was involved in fatty acid and lipid biosynthesis and degradation, consistent with the results of Bi et al (2016) who showed that lipid metabolism is enriched in cortical infection stage of gall development.

The development of this *P. brassicae* - specific database is thus a useful tool for integrating transcriptomics data, gene and metabolic networks and to gain further information than just a list of over- or under-expressed genes between experimental conditions.

Moreover, ClubrootCyc also allowed us to access new pathways and to complete annotation data. For instance, a complete pathway for the spermidine biosynthesis was predicted in eH *P. brassicae* (Figure S6). Particularly, the three enzymes (EC 2.7.2.11, EC 1.2.1.41, and EC 2.6.1.13) required for conversion of glutamate to ornithine, were present in the eH genome although they were not found in the genomes described by Rolfe et al. (2016) or by Schwelm et al. (2015).

Ornithine is converted into putrescine, a precursor of the polyamine spermidine. The roles of polyamines synthesised by plants are well documented. They are involved in fundamental cellular processes, such as chromatin organization, cell proliferation, differentiation, and programmed cell death (Thomas and Thomas, 2001; Bais and Ravishankar, 2002), but also in adaptive responses to abiotic (Bouchereau et al., 1999; Urano et al., 2003; Kuznetsov et al., 2006) and biotic stresses (Walters, 2003). Polyamines have been found being accumulated in the galls caused by *P. brassicae* (Walters and Shuttleton, 1985). In turnip (*Brassica rapa var. rapa)*, the levels of putrescine, spermidine and spermine were reported to be higher in the roots infected with clubroot than in the non-infected normal roots (Walters 2003). In cabbage, broccoli and Komatsuna clubroot galls, the levels of putrescine, spermidine and spermine also increased after infection with *P. brassicae* (Hamana et al., 2015). The high level of polyamines measured in the roots may be contributed by the host plant, as reported by Cao et al. (2008) who showed the up-regulation of the *B. napus* polyamine biosynthetic enzyme spermidine synthase in response to *P. brassicae* infection. However, it could also be due to *P. brassicae* since it is difficult to distinguish between metabolites of plant or pathogen origin. Polyamine biosynthesis could be vital to the survival and pathogenesis of *P. brassicae* as described in other pathogens, such as bacteria (Shah et al., 2011), fungi (Valdes-Santiago et al., 2012) and protists (Rhee et al., 2007).

The chorismate/Shikimate pathway, which was predicted by Schwelm et al. (2015) but lacking the chorismate mutase (EC 5.4.99.5), was found to be complete in the eH *P. brassicae* genome suggesting that this pathway is operational in *P. brassicae*. This pathway is known in plants, bacteria, fungi, and apicomplexan parasites and is central for the biosynthesis of carbocyclic aromatic compounds.

Chorismate is the common branchpoint for the production of these metabolites, and some intermediates in these pathways also have important physiological relevance. For example, salicylate, which is derived from chorismate, is involved in plant defense against *P. brassicae* (Lemarié et al., 2015). Anthranilate, another derivative of chorismate, is the precursor of the *Pseudomonas* quinolone signal, which regulates numerous virulence factors in *Pseudomonas aeruginosa* (Palmer et al., 2013). Furthermore, the chorismate mutase enzyme of this pathway, newly identified in this study in the eH genome, was described as being involved in gall formation in *Meloidogyne javanica* (Doyle and Lambert, 2003) and in *Ustilago maydis* (Djamei et al., 2011). The potential role of this enzyme in *P. brassicae* pathogenicity would require functional studies.

ClubrootCyc also predicted the L-Dopachrome biosynthesis pathway, which is involved in melanin synthesis. Melanin is a broad term for a group of natural pigments found in most organisms and localized in cell walls of different structures of fungi (Toledo et al., 2017). They represent well-known virulence factors for several pathogenic fungi (Jacobson, 2000) and for bacteria (Nosanchuk and Casadevall, 2003).

Metabolic pathway prediction in *P. brassicae* is still, however, incomplete and poorly annotated because there is still not enough information in the reference databases. To make further progress in metabolic network reconstruction and modelling, appropriate gap-filling methods are required. For this, software is being developed to identify and add lacking pathways, such as Meneco, a tool dedicated to the topological gap-filling of genome-scale draft metabolic networks (Prigent et al., 2017).

## 4. Conclusion

In this study, we investigated the mitochondrial sequence of *P. brassicae,* which can now be used for phylogenetic analyses in the supergroup Rhizaria. We also improved the metabolic pathway descriptions for this protist. This allowed the biosynthesis pathways of some specific compounds that could play a role in infection to be clarified. By combining genomic analyses with metabolic network reconstruction in *P. brassicae,* the present study provides useful post-genomics tools devoted to this protist. ClubrootCyc is freely accessible at https://pathway-tools.toulouse.inra.fr/CLUBROOT and can be updated with newly published results. It provides a way to easily visualize omics data and represent metabolic pathways potentially involved in particular experimental conditions of interest. This will help improve our understanding of the infection processes involved in the diseases caused the pathogens of this family by targeting experiments to assess the function of, as yet, unknown enzymes or transporters.

## Acknowledgments

We thank the BRC *BrACySol* (INRA Rennes, France) for providing the seeds. We thank Leigh Gebbie for revising the English written style of the manuscript.

## References

Albataineh MT, Kadosh D (2016) Regulatory roles of phosphorylation in model and pathogenic fungi. Medical Mycology 54, 333–352.

Alborzi SZ, Devignes MD, Ritchie DW (2017) ECDomainMiner: discovering hidden associations between enzyme commission numbers and Pfam domains. BMC Bioinformatics 18, 107.

Al-Khodor S, Price CT, Kalia A, Abu Kwaik Y (2010) Ankyrin-repeat containing proteins of microbes: a conserved structure with functional diversity. Trends in Microbiology 18, 132–139.

Altschul SF, Madden TL, Schäffer AA, Zhang J, Zhang Z, Miller W, Lipman DJ (1997) Gapped BLAST and PSI-BLAST: a new generation of protein database search programs. Nucleic Acids Research 25, 3389–3402.

Ashburner M, Ball CA, Blake JA, Botstein D, Butler H, Cherry JM, Davis AP, Dolinski K, Dwight SS, Eppig JT, Harris MA, Hill DP, Issel-Tarver L, Kasarskis A, Lewis S, Matese JC, Richardson JE, Ringwald M, Rubin GM, Sherlock G (2000) Gene ontology: tool for the unification of biology. The Gene Ontology Consortium Nature Genetics 25, 25–29.

Aurrecoechea C, Barreto A, Basenko E Y, Brestelli J, Brunk BP, Cade S, Crouch K, Doherty R, Falke D, Fischer S, Gajria B, Harb OS, Heiges M, Hertz-Fowler C, Hu S, Iodice J, Kissinger JC, Lawrence C, Li W, Pinney DF, Pulman JA, Roos DS, Shanmugasundram A, Silva-Franco F, Steinbiss S, Stoeckert CJ, Spruill D, Wang H, Warrenfeltz S, Zheng J (2017) EuPathDB: the eukaryotic pathogen genomics database resource. Nucleic Acids Research 45, D581–D591.

Bais HP, Ravishankar GA (2002) Role of polyamines in the ontogeny of plants and their biotechnological applications. Plant Cell, Tissue and Organ Culture 69, 1–34.

Baxter L, Tripathy S, Ishaque N, Boot N, Cabral A, Kemen E, Thines M, Ah-Fong A, Anderson R, Badejoko W, Bittner-Eddy P, Boore JL, Chibucos MC, Coates M, Dehal P, Delehaunty K, Dong S, Downton P, Dumas B, Fabro G, Fronick C, Fuerstenberg SI, Fulton L, Gaulin E, Govers F, Hughes L, Humphray S, Jiang RHY, Judelson H, Kamoun S, Kyung K, Meijer H, Minx P, Morris P, Nelson J, Phuntumart V, Qutob D, Rehmany A, Rougon-Cardoso A, Ryden P, Torto-Alalibo T, Studholme D, Wang Y, Win J, Wood J, Clifton SW, Rogers J, Van den Ackerveken G, Jones JDG, McDowell JM, Beynon J, Tyler BM (2010) Signatures of adaptation to obligate biotrophy in the Hyaloperonospora arabidopsidis genome. Science 330, 1549–1551.

Becht P, Vollmeister E, Feldbrügge M (2005) Role for RNA-binding proteins implicated in pathogenic development of Ustilago maydis. Eukaryotic Cell 4, 121–133.

Bi K, He Z, Gao Z, Zhao Y, Fu Y, Cheng J, Xie J, Jiang D, Chen T (2016) Integrated omics study of lipid droplets from Plasmodiophora brassicae. Scientific Reports 6, 36965.

Bouchereau A, Aziz A, Larher F, Martin-Tanguy J (1999) Polyamines and environmental challenges: recent development. Plant Science 140, 103–125.

Brodmann D, Schuller A, Ludwig-Muller J, Aeschbacher RA, Wiemken A, Boller T, Wingler A (2002) Induction of trehalase in Arabidopsis plants infected with the trehalose-producing pathogen Plasmodiophora brassicae. Molecular Plant-Microbe Interactions 15, 693–700.

Bulman SR, Braselton JP (2014) Rhizaria: Phytomyxea. In: Mc Laughlin DJ, Spatafora JW (eds), The Mycota VII, 2^nd^ ed. Systematics and Evolution, Part A. Springer-Verlag. Berlin-Heidelberg, Germany. pp. 99–112.

Burger G, Gray MW, Forget L, Lang BF (2013) Strikingly bacteria-like and gene-rich mitochondrial genomes throughout Jakobid protists. Genome Biology and Evolution 5, 418–438.

Burki F, Kudryavtsev A, Matz MV, Aglyamova GV, Bulman S, Fiers M, Keeling PJ, Pawlowski J (2010) Evolution of Rhizaria: new insights from phylogenomic analysis of uncultivated protists. BMC Evolutionary Biology 10, 377.

Cao T, Srivastava S, Rahman MH, Kav NNV, Hotte N, Deyholos MK, Strelkov SE (2008) Proteome-level changes in the roots of Brassica napus as a result of Plasmodiophora brassicae infection. Plant Science 174, 97–115.

Caspi R, Billington R, Ferrer L, Foerster H, Fulcher CA, Keseler IM, Kothari A, Krummenacker M, Latendresse M, Mueller LA, Ong Q, Paley S, Subhraveti P, Weaver DS, Karp PD (2016) The MetaCyc database of metabolic pathways and enzymes and the BioCyc collection of pathway/genome databases. Nucleic Acids Research 44, D471–D480.

Conesa A, Götz S, Garcia-Gomez JM, Terol J, Talon M, Robles M (2005) Blast2GO: a universal tool for annotation, visualization and analysis in functional genomics research. Bioinformatics 21, 3674–3676.

Curtis BA, Tanifuji G, Burki F, Gruber A, Irimia M, Maruyama S, Arias MC, Ball SG, Gile GH, Hirakawa Y, Hopkins JF, Kuo A, Rensing SA, Schmutz J, Symeonidi A, Elias M, Eveleigh RJ, Herman EK, Klute MJ, Nakayama T, Oborník M, Reyes-Prieto A, Armbrust EV, Aves SJ, Beiko RG, Coutinho P, Dacks JB, Durnford DG, Fast NM, Green BR, Grisdale CJ, Hempel F, Henrissat B, Höppner MP, Ishida K, Kim E, Kořený L, Kroth PG, Liu Y, Malik SB, Maier UG, McRose D, Mock T, Neilson JA, Onodera NT, Poole AM, Pritham EJ, Richards TA, Rocap G, Roy SW, Sarai C, Schaack S, Shirato S, Slamovits CH, Spencer DF, Suzuki S, Worden AZ, Zauner S, Barry K, Bell C, Bharti AK, Crow JA, Grimwood J, Kramer R, Lindquist E, Lucas S, Salamov A, McFadden GI, Lane CE, Keeling PJ, Gray MW, Grigoriev IV, Archibald JM (2012) Algal genomes reveal evolutionary mosaicism and the fate of nucleomorphs. Nature 492, 59–65.

Devos S, Laukens K, Deckers P, Van Der Straeten D, Beeckman T, Inzé D, Van Onckelen H, Witters E, Prinsen E (2006) A hormone and proteome approach to picturing the initial metabolic events during Plasmodiophora brassicae infection on Arabidopsis. Molecular Plant-Microbe Interactions 19, 1431–1443.

Diederichsen E, Frauen M, Linders EG, Hatakeyama K, Hirai M (2009) Status and perspectives of clubroot resistance breeding in crucifer crops. Journal of Plant Growth Regulation 28, 265–281.

Dierckxsens N, Mardulyn P, Smits G (2017) NOVOPlasty: de novo assembly of organelle genomes from whole genome data. Nucleic Acids Research 45, e18.

Dixon GR (2009) The occurrence and economic impact of Plasmodiophora brassicae and clubroot disease Journal of Plant Growth Regulation 28, 194–202.

Djamei A, Schipper K, Rabe F, Ghosh A, Vincon V, Kahnt J, Osorio S, Tohge T, Fernie AR, Feussner I, Feussner K, Meinicke P, Stierhof YD, Schwarz H, Macek B, Mann M, Kahmann R (2011) Metabolic priming by a secreted fungal effector. Nature 478, 395–398.

Doyle EA, Lambert KN (2003) Meloidogyne javanica chorismate mutase 1 alters plant cell development. Molecular Plant-Microbe Interactions 16, 123–131.

Fabregat A, Sidiropoulos K, Garapati P, Gillespie M, Hausmann K, Haw R, Jassal B, Jupe S, Korninger F, McKay S, Matthews L, May B, Milacic M, Rothfels K, Shamovsky V, Webber M, Weiser J, Williams M, Wu G, Stein L, Hermjakob H, D’Eustachio P (2016) The Reactome pathway knowledgebase. Nucleic Acids Research 44, D481–D487.

Fähling M, Graf H, Siemens J (2003) Pathotype separation of Plasmodiophora brassicae by the host plant. Journal of Phytopathology – Phytopathologische Zeitschrift 151, 425–430.

Feng J, Hwang R, Hwang SF, Strelkov SE, Gossen BD, Zhou QX, Peng G (2010) Molecular characterization of a serine protease Pro1 from Plasmodiophora brassicae that stimulates resting spore germination. Molecular Plant Pathology 11, 503–512.

Feng J, Hwang SF, Strelkov SE (2013) Assessment of gene expression profiles in primary and secondary zoospores of Plasmodiophora brassicae by dot blot and real-time PCR. Microbiological Research 168, 518–524.

Foissac S, Gouzy J, Rombauts S, Mathé C, Amselem J, Sterck L, Van de Peer Y, Rouzé P, Schiex T (2008) Genome Annotation in Plants and Fungi: EuGène as a Model Platform. Current Bioinformatics 3, 87–97.

Glöckner G, Hulsmann N, Schleicher M, Noegel AA, Eichinger L, Gallinger C, Pawlowski J, Sierra R, Euteneuer U, Pillet L, Moustafa A, Platzer M, Groth M, Szafranski K, Schliwa M (2014). The genome of the Foraminiferan Reticulomyxa filosa. Current Biology 24, 11–18.

Gravot A, Grillet L, Wagner G, Jubault M, Lariagon C, Baron C, Deleu C, Delourme R, Bouchereau A, Manzanares-Dauleux MJ (2011) Genetic and physiological analysis of the relationship between partial resistance to clubroot and tolerance to trehalose in Arabidopsis thaliana. New Phytologist 191, 1083–1094.

Gravot A, Deleu C, Wagner G, Lariagon C, Lugan R, Todd C, Wendehenne D, Delourme R, Bouchereau A, Manzanares-Dauleux MJ (2012) Arginase induction represses gall development during clubroot infection in Arabidopsis. Plant and Cell Physiology 53, 901–911.

Gutiérrez P, Bulman S, Alzate J, Ortíz MC, Marín M (2014) Mitochondrial genome sequence of the potato powdery scab pathogen. Mitochondrial DNA 27, 58–59.

Hahn C, Bachmann L, Chevreux B (2013) Reconstructing mitochondrial genomes directly from genomic next-generation sequencing reads - a baiting and iterative mapping approach. Nucleic Acids Research 41, e129.

Hamana K, Hayashi H, Niitsu M (2015) Polyamines in different organs of Brassica crop plants with or without clubroot disease. Plant Production Science 18, 476–480.

Ingram DS, Tommerup IC (1972) The life history of Plasmodiophora brassicae

Woron. Proceedings of the Royal Society B: Biological Sciences 180, 103–112.

Jacobson ES (2000) Pathogenic roles for fungal melanins. Clinical Microbiology Reviews 13, 708–717.

Johnson LN (2009) The regulation of protein phosphorylation. Biochemical Society Transactions 37, 627–641.

Kanehisa M, Furumichi M, Tanabe M, Sato Y, Morishima K (2017) KEGG: new perspectives on genomes, pathways, diseases and drugs. Nucleic Acids Research 45, D353–D361.

Karp PD, Latendresse M, Caspi R (2011) The pathway tools pathway prediction algorithm. Standards in Genomic Sciences 5, 424–429.

Kemen E, Gardiner A, Schultz-Larsen T, Kemen AC, Balmuth AL, Robert-Seilaniantz A, Bailey K, Holub E, Studholme DJ, MacLean D, Jones JDG (2011) Gene gain and loss during evolution of obligate parasitism in the white rust pathogen of Arabidopsis thaliana. PLoS Biology 9, e1001094.

Krogh A, Larsson B, von Heijne G, Sonnhammer ELL (2001) Predicting transmembrane protein topology with a hidden markov model: application to complete genomes. Journal of Molecular Biology 305, 567–580.

Krzywinski M, Schein J, Birol İ, Connors J, Gascoyne R, Horsman D, Jones SJ, Marra MA (2009) Circos: an information aesthetic for comparative genomics. Genome Research. 19, 1639–1645.

Kuznetsov VV, Radyukina NL, Shevyakova NI (2006) Polyamines and stress: biological role, metabolism, and regulation. Russian Journal of Plant Physiology 53, 583–604.

Lemarié S, Robert-Seilaniantz A, Lariagon C, Lemoine J, Marnet N, Jubault M, Manzanares-Dauleux MJ, Gravot A (2015) Both the jasmonic acid and the salicylic acid pathways contribute to resistance to the biotrophic clubroot agent Plasmodiophora brassicae in Arabidopsis. Plant and cell Physiology 56, 2158–2168.

Ludwig-Müller J (2009a) Glucosinolates and the clubroot disease: defense compounds or auxin precursors? Phytochemistry Reviews 8, 135–148.

Ludwig-Müller J (2009b) Plant defence - what can we learn from clubroots? Australasian Plant Pathology 38, 318–324.

Ludwig-Müller J, Jülke S, Geiss K, Richter F, Mithöfer A, Šola I, Rusak G, Keenan S, Bulman S (2015) A novel methyltransferase from the intracellular pathogen Plasmodiophora brassicae methylates salicylic acid. Molecular Plant Pathology 16, 349–364.

Malinowski R, Novak O, Borhan MH, Spichal L, Strnad M, Stephen A. Rolfe SA (2016) The role of cytokinins in clubroot disease. European Journal of Plant Pathology 145, 543–557.

Manzanares-Dauleux MJ, Barret P, Thomas G (2000) Development of a pathotype specific SCAR marker in Plasmodiophora brassicae. European Journal of Plant Pathology 106, 781–787.

Molmeret M, Horn M, Wagner M, Santic M, Abu Kwaik A (2005) Amoebae as training grounds for intracellular bacterial pathogens. Applied Environmental Microbiology 71, 20–28.

Nosanchuk JD, Casadevall A (2003) The contribution of melanin to microbial pathogenesis. Cellular Microbiology 5, 203–223.

Palmer GC, Jorth PA, Whiteley M (2013) The role of two Pseudomonas aeruginosa anthranilate synthases in tryptophan and quorum signal production. Microbiology 159, 959–969.

Petersen TN, Brunak S, von Heijne G, Nielsen H (2011) SignalP 4.0: discriminating signal peptides from transmembrane regions. Nature Methods 8, 785– 786.

Prigent S, Frioux C, Dittami SM, Thiele S, Larhlimi A, Collet G, Gutknecht F, Got J, Eveillard D, Bourdon J, Plewniak F, Tonon T, Siegel A (2017) Meneco, a topology-based gap-filling tool applicable to degraded genome-wide metabolic networks. PLoS Computational Biology 13, e1005276.

Rhee HJ, Kim E-J, Lee JK (2007) Physiological polyamines: simple primordial stress molecules. Journal of Cellular and Molecular Medicine 11, 685–703.

Rizk G, Gouin A, Chikhi R, Lemaitre C (2014) MindTheGap: integrated detection and assembly of short and long insertions. Bioinformatics 30, 3451–3457.

Rolfe SA, Strelkov SE, Links MG, Clarke WE, Robinson SJ, Djavaheri M, Malinowski R, Haddadi P, Kagale S, Parkin IAP, Taheri A, Borhan MH (2016) The compact genome of the plant pathogen Plasmodiophora brassicae is adapted to intracellular interactions with host Brassica spp. BMC Genomics, 17, 272.

Schuller A, Kehr J, Ludwig-Müller J (2014) Laser microdissection coupled to transcriptional profiling of Arabidopsis roots inoculated by Plasmodiophora brassicae indicates a role for brassinosteroids in clubroot formation. Plant and Cell Physiology 55, 392–411.

Schwelm A, Fogelqvist J, Knaust A, Jülke S, Lilja T, Bonilla-Rosso G, Karlsson M, Shevchenko A, Dhandapani V, Choi SR, Kim HG, Park JY, Lim YP, Ludwig-Müller J, Dixelius C (2015) The Plasmodiophora brassicae genome reveals insights in its life cycle and ancestry of chitin synthases. Scientific Reports 5, 11153.

Schwelm A, Dixelius C, Ludwig-Müller J (2016) New kid on the block – the clubroot pathogen genome moves the plasmodiophorids into the genomic era. European Journal of Plant Pathology 145, 531–542.

Shah P, Nanduri B, Swiatlo E, Ma Y, Pendarvis K (2011) Polyamine biosynthesis and transport mechanisms are crucial for fitness and pathogenesis of Streptococcus pneumoniae. Microbiology 157, 504–515.

Siemens J, Keller I, Sarx J, Kunz S, Schuller A, Nagel W, Schmülling T, Parniske M, Ludwig-Müller J (2006) Transcriptome analysis of Arabidopsis clubroots Indicate a key role for cytokinins in disease development. Molecular Plant-Microbe Interactions 19, 480–494.

Somé A, Manzanares MJ, Laurens F, Baron F, Thomas G, Rouxel F (1996) Variation for virulence on Brassica napus L amongst Plasmodiophora brassicae collections from France and derived single-spore isolates. Plant Pathology 45, 432– 439.

Spanu PD, Abbott JC, Amselem J, Burgis TA, Soanes DM, Stüber K, Ver Loren van Themaat E, Brown JK, Butcher SA, Gurr SJ, Lebrun MH, Ridout CJ, Schulze-Lefert P, Talbot NJ, Ahmadinejad N, Ametz C, Barton GR, Benjdia M, Bidzinski P, Bindschedler LV, Both M, Brewer MT, Cadle-Davidson L, Cadle-Davidson MM, Collemare J, Cramer R, Frenkel O, Godfrey D, Harriman J, Hoede C, King BC, Klages S, Kleemann J, Knoll D, Koti PS, Kreplak J, López-Ruiz FJ, Lu X, Maekawa T, Mahanil S, Micali C, Milgroom MG, Montana G, Noir S, O’Connell RJ, Oberhaensli S, Parlange F, Pedersen C, Quesneville H, Reinhardt R, Rott M, Sacristán S, Schmidt SM, Schön M, Skamnioti P, Sommer H, Stephens A, Takahara H, Thordal-Christensen H, Vigouroux M, Wessling R, Wicker T, Panstruga R (2010) Genome expansion and gene loss in powdery mildew fungi reveal tradeoffs in extreme parasitism. Science 330, 1543–1546.

Supek F, Bosnjak M, Skunca N, Smuc T (2011) REVIGO summarizes and visualizes long lists of Gene Ontology terms. PLoS One 6, e21800.

Tanifuji G, Archibald JM, Hashimoto T (2016) Comparative genomics of mitochondria in chlorarachniophyte algae: endosymbiotic gene transfer and organellar genome dynamics. Scientific Reports 6, 21016.

The Gene Ontology Consortium (2017) Expansion of the Gene Ontology knowledgebase and resources. Nucleic Acids Research 45, D331–D338.

The UniProt Consortium (2017) UniProt: the universal protein knowledgebase. Nucleic Acids Research 45, D158–D169.

Thomas T, Thomas TJ (2001) Polyamines in cell growth and cell death: molecular mechanisms and therapeutic applications. Cellular and Molecular Life Sciences 58, 244–258.

Toledo AV, Franco MEE, Lopez SMY, Troncozo MI, Saparrat MCN, Balatti PA (2017) Melanins in fungi: types, localization and putative biological roles. Physiological and Molecular Plant Pathology 99, 2–6.

Turra D, Segorbe D, Di Pietro A (2014) Protein kinases in plant-pathogenic fungi: conserved regulators of infection. Annual Review of Phytopathology 52, 267–288.

Urano K, Yoshiba Y, Nanjo T, Igarashi Y, Seki M, Sekiguchi F, Yamaguchi-Shinozaki K, Shinozaki K (2003) Characterization of Arabidopsis genes involved in biosynthesis of polyamines in abiotic stress responses and developmental stages. Plant, Cell & Environment 26, 1917–1926.

Valdes-Santiago L, Cervantes-Chavez JA, Leon-Ramırez CG, Ruiz-Herrera J (2012) Polyamine metabolism in fungi with emphasis on phytopathogenic species. Journal of Amino Acids 2012, 13 pages.

Wagner G, Charton S, Lariagon C, Laperche A, Lugan R, Hopkins J, Frendo P, Bouchereau A, Delourme R, Gravot A, Manzanares-Dauleux MJ (2012) Metabotyping: A new approach to investigate rapeseed (Brassica napus L.) genetic diversity in the metabolic response to clubroot infection. Molecular Plant-Microbe Interactions 25, 1478–1491.

Walters DR (2003) Polyamines and plant disease. Phytochemistry 64, 97–107.

Walters DR, Shuttleton MA (1985) Polyamines in the roots of turnip infected with Plasmodiophora brassicae Wor. New Phytologist 100, 209–214.

Yin Y, Mao X, Yang J, Chen X, Mao F, Xu Y (2012) dbCAN: a web resource for automated carbohydrate-active enzyme annotation. Nucleic Acids Research 40, W445–W451.

